# From lake to estuary, the tale of two waters. A study of aquatic continuum biogeochemistry

**DOI:** 10.1101/184101

**Authors:** Paul Julian, Todd Z. Osborne

## Abstract

The balance of fresh and saline water is essential to estuarine ecosystem function. Along the fresh-brackish-saline water gradient within the C-43 canal/Caloosahatchee River estuary (CRE) the quantity, timing and distribution of water and associated water quality significantly influences ecosystem function. Long-term trends of water quality and quantity were assessed from Lake Okeechobee to the CRE between May 1978 to April 2016. Significant changes to monthly flow volumes were detected between the lake and the estuary which correspond to changes in upstream management. and climatic events. Across the 37-year period total phosphorus (TP) flow-weighted mean (FWM) concentration significantly increased at the lake, meanwhile total nitrogen (TN) FMW concentrations significantly declined at both the lake and estuary headwaters. Between May 1999 and April 2016, TN, TP and total organic carbon (TOC), ortho-P and ammonium conditions were assessed within the estuary at several monitoring locations. Generally nutrient concentrations decreased from upstream to downstream with shifts in TN:TP from values >20 in the freshwater portion, ~20 in the estuarine portion and <20 in the marine portion indicating a spatial shift in nutrient limitations along the continuum. Aquatic productivity analysis suggests that the estuary is net heterotrophic with productivity being negatively influenced by TP, TN and TOC likely due to a combination of effects including shading by high color dissolved organic matter. We conclude that rainfall patterns, land use and the resulting discharges of run-off drives the ecology of the C-43/CRE aquatic continuum and associated biogeochemistry rather than water management associated with Lake Okeechobee.

## Introduction

The aquatic continuum concept (ACC) originated from an ecohydrological concept to understand the transport of material between landscape level patches (i.e. ecosystems) through the aquatic-terrestrial continuum (Jenerette and Lal 2005). Within the literature the ACC is at times synonymous with the river continuum concept (RCC) developed by (Vannote et al. 1980), which formulated a conceptual model and framework for characterizing lotic (running water) ecosystems to describe the ecology of communities along a river system. Both the ACC and RCC bridge terrestrial and aquatic biogeochemical processes by shared fundamental concepts such as nutrient limitation, ecosystem nutrient retention and controls of nutrient transitions (Grimm et al. 2003). The one important link between the terrestrial and aquatic ecosystem is the flow of water and energy (Abrantes and Sheaves 2010; England and Rosemond 2004; Mendoza–Lera et al. 2012). Sediment, organic carbon and nutrient loading from land journey through the continuum that includes a path through soils to the open ocean and all compartments between (i.e. groundwater, floodplains, rivers, lakes, estuaries, etc.). The progression of materials and energy through these systems act as a succession of filters in which the hydrology, ecology and biogeochemical processing are tightly coupled and act to retain a significant fraction of the nutrients transported through each system (Bouwman et al. 2013). Retention of nutrients along the aquatic continuum not only influences the total amount of nutrients reaching the aquatic end-member (i.e. ocean) but also how the systems modify the stoichiometry and forms of these nutrients en-route (Billen 1993; Ensign and Doyle 2006; Reddy et al. 1999). However, anthropogenic alteration of the ecosystem has changed the hydrology and nutrient stoichiometry and forms along the aquatic continuum.

Primary production forms a critical base to estuarine food webs, strongly influencing oxygen and nutrient dynamics (Caffrey et al. 2013). The magnitude and relative values of gross primary productivity (GPP) and ecosystem respiration (ER) can vary due to flow (stagnant versus flowing conditions), inputs of terrestrial or anthropogenic organic carbon (OC) and nutrients (Odum 1956). Organic matter (OM) in river and estuarine systems are principally derived from terrestrial vegetation and soils, considered recalcitrant and transported conservatively to the ocean. However, several biogeochemical processes can contribute to non-conservative behavior of OM in estuarine ecosystems (Bianchi 2013). Nutrients enter estuaries from either terrestrial sources via run-off, upstream from freshwater rivers and wetlands or tidal exchange of marine waters and in some areas, terrestrial inputs of nutrients have had deleterious effects on estuarine and coastal ecosystems (Bianchi 2013; Jickells 1998). Hydrologic pulsing of freshwater and the associated delivery of OC and nutrients have been observed to influence net aquatic productivity in river, wetland and coastal wetland ecosystems (Gallardo et al. 2012; Maynard et al. 2012; Shen et al. 2015).

Inflow of freshwater to an estuary is a significant landscape process that shapes community structure (Mannino and Montagna 1997). The management of freshwater inflows to estuaries can have profound effects on conditions and ecosystem function (Alber 2002; Kimmerer 2002; Sklar and Browder 1998). Similar to many urbanized coastal areas, south Florida estuaries have been significantly altered (Macauley et al. 2002). Along with the loss of shoreline habitat and function, the draining and channelization of wetlands for the purposes of agriculture and diversion of water for urban development has led to increased wet season flows and decreased dry season flows (Doering and Chamberlain 1999). This change in flow patterns has changed the distribution and abundance of key estuarine indicator species such as seagrasses and oysters (Buzzelli et al. 2015). Oysters are sensitive to salinity and siltation (T. K. Barnes et al. 2007; Soniat et al. 2013) while salinity patterns also influence the composition, distribution and abundance of seagrass habitats in estuarine ecosystems (Doering et al. 2002; Greenawalt-Boswell et al. 2006; Livingston et al. 1998). Unlike other estuaries, artificial connections to Lake Okeechobee create the potential for transport of material to the Caloosahatchee and St. Lucie Estuaries from an expanded drainage area extending to the chain of lakes near Orlando, a distance of over 300 km. In combination with alterations to the local watersheds of both systems (Doering et al. 2006; Sun et al. 2016; Wan et al. 2014), this connection has resulted in significant alteration to the timing, distribution and volume of water entering these estuaries.

The completion of the Lake Okeechobee levy in 1937 brought the loss of connectivity with the Florida Everglades and reduced its total area by 50% (Steinman et al. 2002) and ultimately disrupting the aquatic continuum. Loss of the Everglades connection also resulted in the loss of significant amounts of water storage and simultaneously increased nutrient loading to the Caloosahatchee and St. Lucie estuaries. While much attention has been directed at Florida Bay, the coral reefs of the Florida Keys and the Everglades wetlands downstream of Lake Okeechobee (Porter and Porter 2002), the conditions within the Caloosahatchee and St. Lucie Estuaries are dictated by base flows, local basin stormwater runoff and regulatory releases from Lake Okeechobee (Doering and Chamberlain 1999; Chamberlain and Hayward 1996). Two major problems for the Caloosahatchee River Estuary (CRE) are excessive nutrient loading and high frequency and duration of undesirable salinity ranges within the estuary (T. Barnes 2005; South Florida Water Management District et al. 2009).

This study had three objectives to contrast the ecological and biogeochemical processes across the aquatic continuum. The first objective was to evaluate changes in water quality and quantity of freshwater flows into the CRE with the hypothesis that upstream water management and localized climatic conditions have increased flows entering the estuary during the baseline period (May 1^st^ 1978 – April 30^th^ 2016) resulting in degraded water quality entering the estuary and increasing long-term trends. The second objective was to investigate water quality characteristics and limiting nutrients testing the hypothesis that the estuary is a “right-side up” with limiting nutrients being supplied by up-stream rather than “upside down” estuaries with limiting nutrients being supplied by marine or ocean sources (Childers et al. 2006a; Walters et al. 1988). The third objective was to evaluate how freshwater quantity and quality influences aquatic productivity in the CRE testing the hypothesis that freshwater flow positively influences productivity at freshwater sites and negatively influence productivity at estuarine or marine sites.

## Methods

### Study Area

The Caloosahatchee River and its estuary are located on the lower west coast of Florida, USA (Fig. 1). The historic Caloosahatchee River was originally shallow, meandering with its headwaters located approximately 100 km inland and tidally influenced up to approximately 70 km upstream. However, to accommodate navigation, flood-control and land-reclamation needs the river was extended eastward and connected to Lake Okeechobee in late 1800’s and early 1900’s by what is now called the C-43 canal. Smaller secondary canals were constructed to irrigate and drain lands surrounding the river. In addition to connecting the river to Lake Okeechobee three lock-and-dam structures were also installed on the river to control flow and stage height. The final downstream structure, the S-79 regulates flow into the estuary and acts as a salinity barrier (T. Barnes 2005; Doering and Chamberlain 1999).

**Figure 1.**
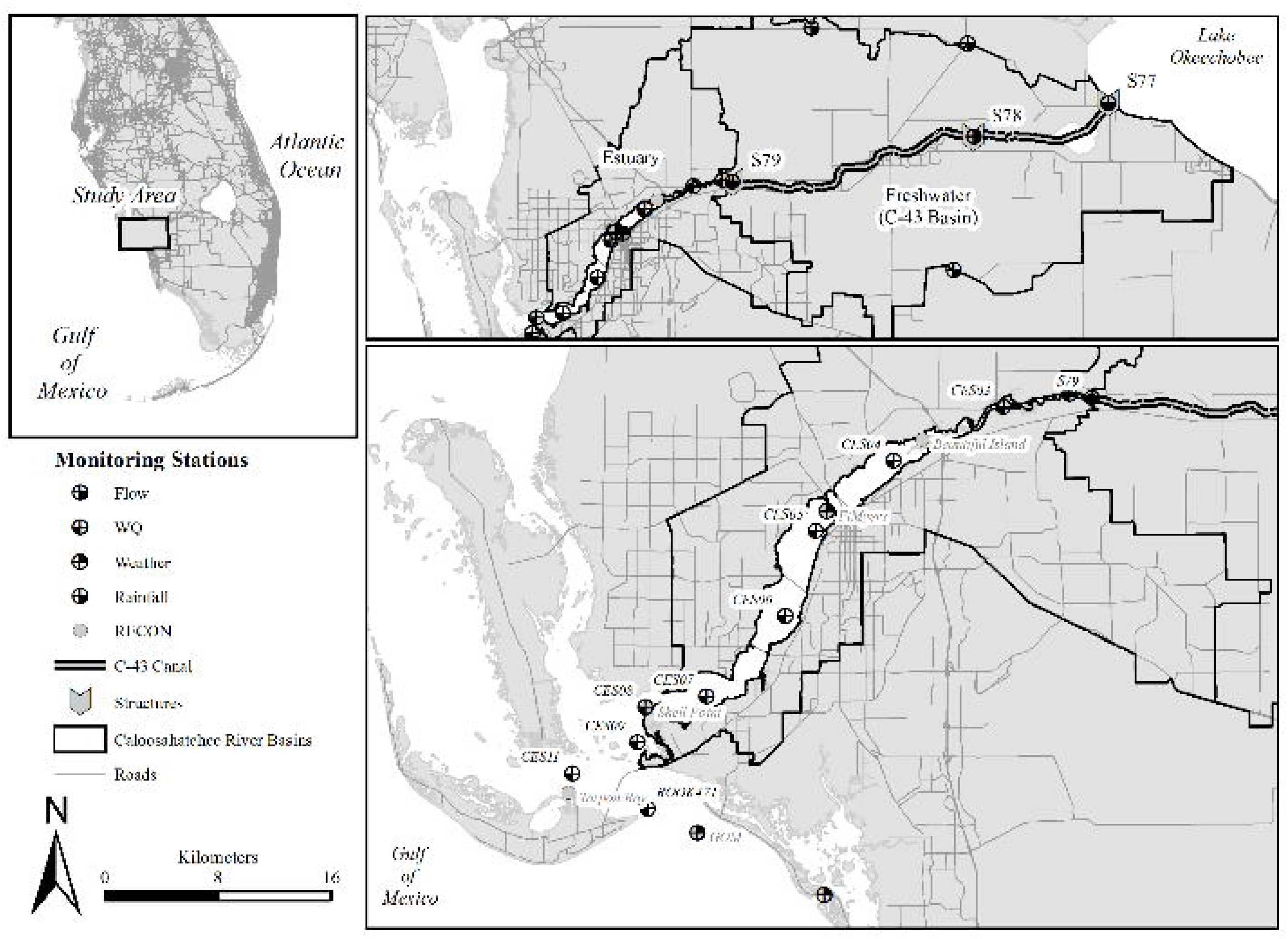
The C-43 and Caloosahatchee River Estuary (CRE) with water quality, River, Estuary and Coastal Observing Network (RECON), flow, weather and rainfall monitoring locations identified. *Upper left insert* identifies study location relative to the rest of peninsula Florida, USA. *Upper right insert* identifies the C-43 canal, C-43 basin and estuary relative to Lake Okeechobee and associated water control structures.

The CRE drains a watershed of approximately 3,450 km^2^ including the C-43 basin (70%) and the tidal basin (30%). Irrigated agricultural lands are among the major land use types in the watershed and impart a significant demand for water during the dry season. The C-43 canal conveys basin runoff and releases for flood control and water supply from Lake Okeechobee to the estuary via the S-79. The pattern and magnitude of flows to the CRE is highly variable due to the subtropical climate receiving approximately 134 cm yr^-1^ of rainfall (historical average from 1965 to 2005) discharges (including flood control and water supply releases) from Lake Okeechobee, and withdrawals for irrigation and water supply for agricultural and urban uses (Qiu and Wan 2013),. Water releases from Lake Okeechobee are made according to a comprehensive regulation schedule used by the U.S. Army Corps of Engineers (USACE) to manage the lake water levels for these diverse and often competing uses (U.S. Army Corps of Engineers and South Florida Water Management District 2010).

### Data Sources

Water quality and hydrologic data were retrieved from the South Florida Water Management District (SFWMD) online database (DBHYDRO; <u>www.sfwmd.gov/dbhydro</u>) for sites within the Caloosahatchee and the C-43 canals (Fig. 1). Water quality sites were grouped by their location within the CRE, and historic salinity values (Supplemental Fig. 1); sites S-79, CES02 and CES03 were classified as freshwater, CES04 to CES09 were classified as estuarine and CES11 and ROOK471 were classified as marine (Fig. 1). The period of record (POR) for flow data spanned the period between water year (WY) 1979 and 2016 (May 1, 1978 – April 30, 2016). Estuarine water quality data was slightly more limited, covering WY2000 to 2016. Water quality parameters used in this study include total phosphorus (TP), orthophosphate (OPO4), total nitrogen (TN), ammonia (NH4) and nitrate-nitrite (NOx) and total organic carbon (TOC). Total nitrogen was either directly measured or calculated from the sum of nitrate-nitrite and total kjeldahl nitrogen. Summary of analytical methods used for each parameter can be found in Table 1.

**Table 1.**
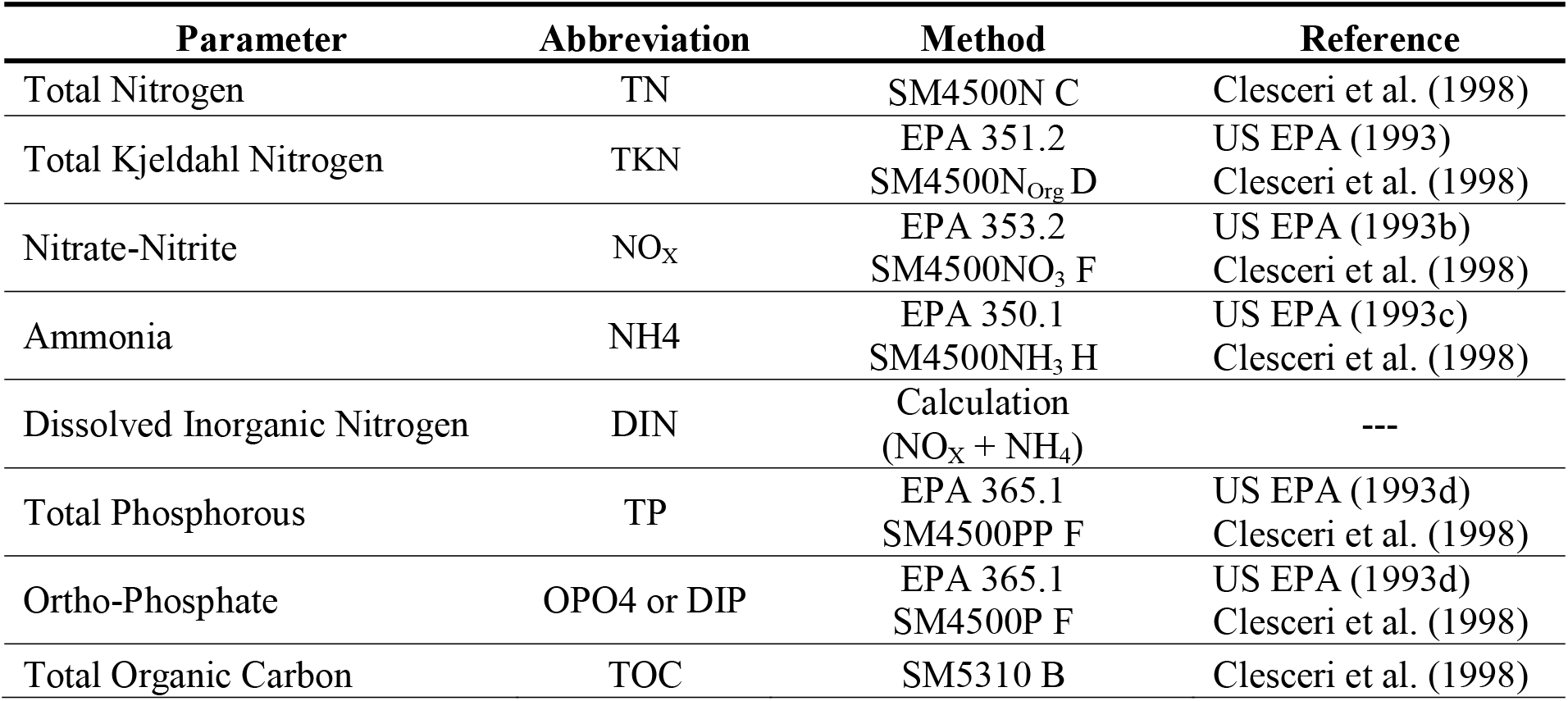
Summary of parameters and analytical methods used for this study.

| Parameter | Abbreviation | Method | Reference |
| --- | --- | --- | --- |
| Total Nitrogen | TN | SM4500N C | Clesceri et al. (1998) |
| Total Kjeldahl Nitrogen | TKN | EPA 351.2<br>SM4500N<sub>Org</sub> D | US EPA (1993)<br>Clesceri et al. (1998) |
| Nitrate-Nitrite | NO<sub>x</sub> | EPA 353.2<br>SM4500NO<sub>3</sub> F | US EPA (1993b)<br>Clesceri et al. (1998) |
| Ammonia | NH<sub>4</sub> | EPA 350.1<br>SM4500NH<sub>3</sub> H | US EPA (1993c)<br>Clesceri et al. (1998) |
| Dissolved Inorganic Nitrogen | DIN | Calculation<br>(NO<sub>x</sub> + NH<sub>4</sub>) | --- |
| Total Phosphorous | TP | EPA 365.1<br>SM4500PP F | US EPA (1993d)<br>Clesceri et al. (1998) |
| Ortho-Phosphate | OPO<sub>4</sub> or DIP | EPA 365.1<br>SM4500P F | US EPA (1993d)<br>Clesceri et al. (1998) |
| Total Organic Carbon | TOC | SM5310 B | Clesceri et al. (1998) |

Water quality data were screened based on laboratory qualifier codes, consistent with FDEP’s quality assurance rule (Florida Administrative Code 2008). Any datum associated with a fatal qualifier indicating a potential data quality problem was removed from the analysis. Additional considerations in the handling of analytical data were the accuracy and sensitivity of the laboratory method used. For purposes of data analysis and summary statistics, data reported as less than method detection limit (MDL) were assigned the maximum MDL determined during this study, unless otherwise noted. Conventionally data less-than the MDL are assigned either the MDL or ½ the MDL, however due to the long POR and changes in MDL values during the study the highest MDL was selected to advert potential biases in long-term trend analyses.

More recently, *in situ* data collected at high frequencies are available for key parameters related to estuary function. The River, Estuary and Coastal Observing Network (RECON) are real-time sensor platforms at fixed sites in the Caloosahatchee Estuary, San Carlos Bay, Pine Island Sound and Gulf of Mexico. For the purposes of this study, only sites within the Caloosahatchee Estuary, San Carlos Bay and Gulf of Mexico were considered (Fig. 1) between WY2009 and 2016 (May 1, 2008 and April 30, 2016). The RECON sensor platforms record several biological, chemical and physical parameters and are mounted to pilings at a depth of 1.5 meters below mean lower low water (MLLW). Instruments are deployed and maintained from small boats with a maximum service interval of one to two months (Milbrandt et al. 2016). Chlorophyll, turbidity, conductivity/salinity, temperature, dissolved oxygen and pressure/depth were measured by the WET Labs Water Quality Monitoring instrument package (WQM) and colored (chromophoric) dissolved organic matter (CDOM) was measured by the WET Labs ECO FLS instrument (ECO). A subset of parameters was used in this study including specific conductance/salinity, temperature, dissolved oxygen and CDOM. The Seabird Coastal instruments (WQM, ECO) were calibrated at the factory annually. During service, sensors were cleaned monthly with a nonionic surfactant (i.e. Triton X-100) and distilled water. The protective cage and exterior surfaces were coated with antifouling paint prior to deployment and after 1 year, the exterior was stripped and re-painted. Discrete samples were collected for each monthly service visit using a YSI EXO2 hand-held sonde at the sensor depth to cross validate *in situ* sensor data.

### Data Analysis

Annual total and monthly flows were summarized for S-77, S-79 and C-43 basin for the Caloosahatchee estuary. C-43 basin flow was estimated as the difference between the S-79 and S-77. Monthly and annual Mann-Kendall trend analysis was performed on each flow datasets from WY1979 to 2016 using the ‘mk.test’ function and Pettitt’s test for change point detection using the ‘pettitt.test’ function in the ‘trend’ R-package (Pholert 2016). Segmented regression was applied to cumulative annual flow volume from the S79 and cumulative annual average rainfall across the watershed to determine changes in the flow-rainfall relationship related to freshwater inputs into the estuary using the ‘segmented’ function in the ‘segmented’ R-package (Vito and Muggeo 2003). Hydraulic loading rate (HLR) was calculated for each water control structure by dividing the total annual flow by the area around the C-43 Canal (~334 km^2^) and the CRE (~108 km^2^) for the S-77 and S-79, respectively.

Annual TP and TN loads and flow-weighted mean (FWM) concentrations were computed for the S-77 and S-79 structures, by interpolating water quality concentration daily from grab samples collected at each respective structure during days of observed flow. Daily interpolated water quality concentrations were then multiplied by daily flow and summed for each WY. Annual FWM values were calculated by dividing total annual nutrient load by total annual flow. Mann-Kendall trend analysis was also performed on annual TP and TN FWM concentrations for the S-77 and S-79 structures using the ‘mk.test’ function and change point detection using the ‘ pettitt.test’. Annual nutrient loads were compared to HLR for both the C-43 canal and CRE using spearman rank correlation.

Annual arithmetic mean concentrations were computed for each station within the CRE with greater than six samples per year. Annual mean concentrations were compared between regions (i.e. freshwater, estuary and marine) using the Dunn’s test for multiple comparison for TP, TN, NH4 and OPO4 between WY2000 to 2016 using the ‘dunn.test’ function in the ‘dunn.test’ R-package (Dinno 2015). Longitudinal gradient analysis of annual TP, TN and TOC concentrations along the estuary was conducted using a Thiel-Sen estimate to derive a non-parametric linear model using the ‘zyp.sen’ function in the ‘zyp’ R-package (Bronaugh and Werner 2013) with greater than six sampling locations along the longitudinal distance of the estuary. Kendall trend analysis was performed on annual slope values of the non-parametric linear model for each parameter (i.e. TP, TN and TOC) by WY.

In addition to decadal trend analysis of water flows and quality, more recent high frequency data allowed for an analysis of aquatic productivity. Aquatic productivity was estimated using hourly DO, surface water temperature, salinity, wind speed, air temperature and barometric pressure data measured at sites along the CRE between May 1^st^ 2009 and April 30^th^ 2016 (Fig. 1). Sampling locations at Beautiful Island, Fort Myers and Shell Point are characteristically estuarine while Tarpon Bay and Gulf of Mexico are characteristically marine. Gross primary productivity (GPP), ecosystem respiration (ER) and net aquatic productivity (NAP) was estimated using the DO rate of change method first proposed by (Odum 1956) and employed by others (Cole et al. 2000; Hagerthey et al. 2010; Staehr et al. 2010; E. Thébault and Loreau 2003). Daily NAP, GPP and ER was calculated on the basis of changes in DO concentrations thought to be driven by the rates of photosynthesis, respiration and atmospheric exchange (Odum 1956). Where NAP and ER can be determined directly from the sensor data, GPP is estimated by mass balance. Within a one-hour interval it is assumed that the change in DO is equality to sum of the respiration rate and oxygen diffusion rate minus the rate of photosynthesis (Equation 1).

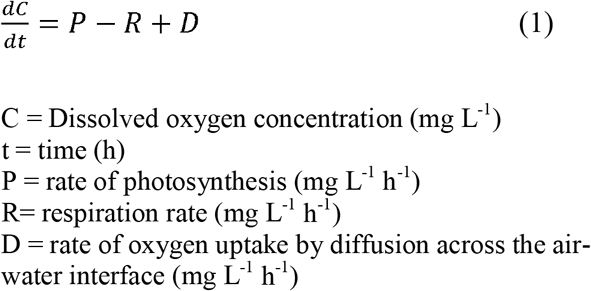

The rate of oxygen uptake by diffusion across the air-water interface (D) is regulated by the difference in O2 in the water column from atmospheric equilibrium and the temperature-dependent gas exchange coefficient for oxygen. Wind produces turbulence in the stationary water bodies, facilitating gas exchange processes driven by wind speed (Equation 2).

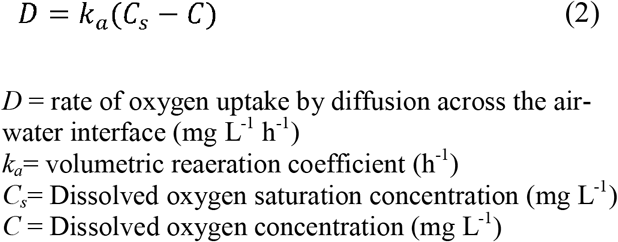

It was assumed that wind speed measured at the Fort Myers weather RECON and the Fort Myers NOAA weather stations would be a reasonable surrogate for the wind conditions for the Beautiful Island and Fort Myers RECON stations. Meanwhile the Gulf of Mexico RECON and the University of South Florida BCP weather stations was used as surrogate for wind conditions at the Gulf of Mexico, Shell Point and Tarpon Bay RECON stations (Fig. 1 and Supplemental Table 1). Volumetric reaeration coefficient (*k_a_*) was calculated using three different functions of wind speed, barometric pressure, air temperature and surface water salinity consistent with methods by (E. Thébault and Loreau 2003) and (Caffrey et al. 2013).

Net aquatic productivity is the difference between photosynthesis and respiration which can be computed by rearranging Equation 1. For each day NAP was calculated by summing hourly diffusion-corrected rates of DO change 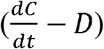 over a 24-hour period, starting and ending at sunrise. Sunrise and sunset were estimated using algorithms developed by NOAA in a modified version of the ‘metab_day’ function in the R package ‘SWMPr’ (Beck 2016). Hourly ER was computed as the mean night-time diffusion corrected rates of DO change, it is assumed that GPP at night is zero (Cole et al. 2000) and the change in DO within an hourly interval during the night is attributed to respiration and diffusion. Assuming that night time ER is equal to daytime ER, daily respiration was calculated by multiplying hourly ER by 24 hours. Hourly photosynthesis was calculated by subtracting ER from diffusion corrected rates of DO change during the daylight period while daily photosynthesis was computed by summing hourly photosynthesis values from sunrise to sunset. Volumetric rates of GPP, ER and NAP were multiplied by depth to yield areal productivity rates (g O_2_ m^-2^ d^-1^) consistent with methods used by others (Caffrey et al. 2013; Cole et al. 2000; Hagerthey et al. 2010; Staehr et al. 2010; E. Thébault and Loreau 2003). Productivity calculations were performed using a modified version of the ‘metabolism’ function in the R package ‘SWMPr’ (Beck 2016).

The ratio between GPP and ER was compared to zero using the Mann-Whitney/Wilcoxon Rank Sum test, if the GPP:ER is significantly less than zero the site would be dominantly heterotrophic meanwhile if the GPP:ER ratio is greater than zero the site would be dominantly autotrophic. Aquatic productivity rates (GPP, ER and NAP) were compared between ecosystems using Kruskal-Wallis rank sum test for the entire POR. River, Estuary and Coastal Observing Network monitoring sites were paired with near-by ambient water quality monitoring locations on days when water quality samples were collected. Spearman’s rank sum correlation was used to compare daily NAP with TP, TN and TOC concentrations, OC:N, OC:P and N:P molar ratios at each RECON-Water Quality paired site. Piecewise regression was used to detect change points in the NAP-TOC, NAP-TP and NAP-TN relationships using the ‘piecewise.regression’ function in the ‘SiZer’ R-package (Sonderegger 2012).

Unless otherwise noted all statistical operations were performed using the base stats R-package. All statistical operations were performed with R© (Ver 3.1.2, R Foundation for Statistical Computing, Vienna Austria). The critical level of significance was set to α = 0.05.

## Results

### C-43 Canal-Caloosahatchee River Estuary Hydrology

Annual flow from the Lake Okeechobee to the C-43 basin via the S-77 ranged from 0.05 to 2.68 km^3^ yr^-1^ and a mean of 0.73 ± 0.11 km^3^ yr^-1^ during the 37-year period between water year 1979 (WY1979) to WY2016 (Fig. 2) with no significant annual trend in annual flow (τ=0.11, ρ=0.35) and a weak increasing trend in monthly total flow volume (τ=0.14, ρ<0.001). A significant change point was detected in monthly flow volume after August 1994 (K=16048, ρ<0.001). Intra- and inter-annual flows entering the C-43 Basin from Lake Okeechobee were highly variable with little seasonality as indicated by monthly (Supplemental Fig. 2) and annual flow volumes (Fig. 2) observed at the S-77 structure across the POR. Furthermore, the POR monthly inter-quantile range is relatively narrow with several monthly values above the 75^th^ quantile.

**Figure 2.**
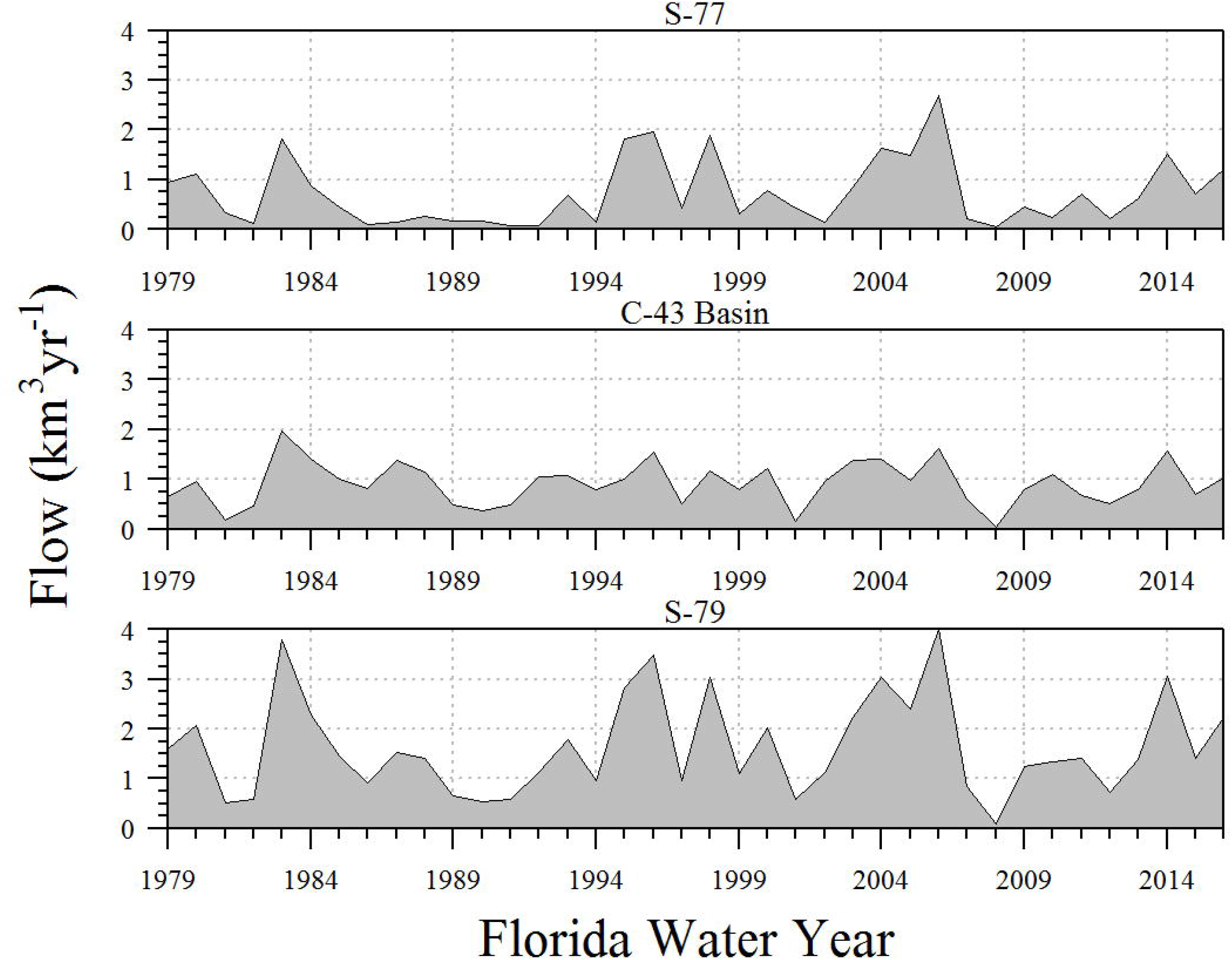
Annual total flow volumes for the C-43 Canal and Caloosahatchee river estuary between WY1979 and WY2016 (May 1, 1978 – April 30, 2016).

The Franklin Lock and Dam (S-79) is the final downstream structure along the C-43 canal and acts as an impediment to saltwater intrusion to the now the C-43 canal (T. Barnes 2005; Buzzelli et al. 2015). Freshwater flows to the CRE via the S-79 ranged from 0.10 to 3.99 km^3^ yr^-1^ during the same period (Fig. 2) with a POR mean 1.64 ± 0.16 km^3^ yr^-1^ with no significant annual trend in total flow volume (τ=0.09, ρ=0.45) and a weak increasing statistically significant monthly (τ=0.09, ρ<0.01) trend in flow volume. Monthly flows from the S-79 exhibit a strong seasonality with peak flows occurring during the August/September time period and low flows occurring between November and April following the characteristic wet season-dry season cycle (Supplemental Fig. 2). Annual flow through the S-79 structure was 174 to 321 percent of the annual flow observed at the S-77, suggesting that in most years, rainfall driven runoff from the C-43 watershed is the dominant source of water to the CRE at S-79.

Due to localized rainfall and runoff within the C-43 basin, annual flow from the basin ranged from 0.04 to 1.96 km^3^ yr^-1^ between WY1979 and 2016 (Fig. 2) with a mean of 0.92 ± 0.07 km^3^ yr^-1^ and no significant annual trend in flow (τ=0.03, ρ=0.82) and a weak increasing trend in monthly total flow volume (τ=0.15, ρ<0.001). Furthermore, a significant change point in the time series of flow volume occurred during June 2007 (K=28610, ρ<0.001), likely a response to a passing tropical storm (Tropical Storm Barry; (Avila 2007) after a prolonged drought period. Monthly basin flow volumes follow the characteristic wet to dry season pattern with the highest flow peaking during August and the lowest between November and April (Supplemental Fig. 2). Furthermore, daily average annual total rainfall across the C-43 basin ranged from 0.82 to 1.94 m yr^-1^ during the POR with a mean of 1.34 ± 0.04 m yr^-1^.

Reviewing the record of flow at the S-79 structure and the C-43 basin rainfall, there were notable changes to the relationship between rainfall and runoff that has occurred throughout the POR (Fig 3). Additionally, the piecewise regression analysis approach identified several statistically significant breakpoints in the cumulative rainfall-discharge relationship for the S-79 structure (R^2^ =1.00, F_(9,28)_ = 2936, ρ<0.001). The breakpoints along the rainfall and runoff relationship for the S-79 correspond to WY1986, 1992, 2007 and 2012, which suggests significant changes in either water management (i.e. S-79 flows), occurrence of extreme climatic events including high rains or droughts or a combination of both. Based on the double mass curve, the periods 1986 to 1992 and 2007 to 2012 periods were relatively dry periods with respect to rainfall (Supplemental Fig. 3) and flow as indicated by the decreases in slope. By comparison, the 1992 to 2007 period was relatively wet.

**Figure 3.**
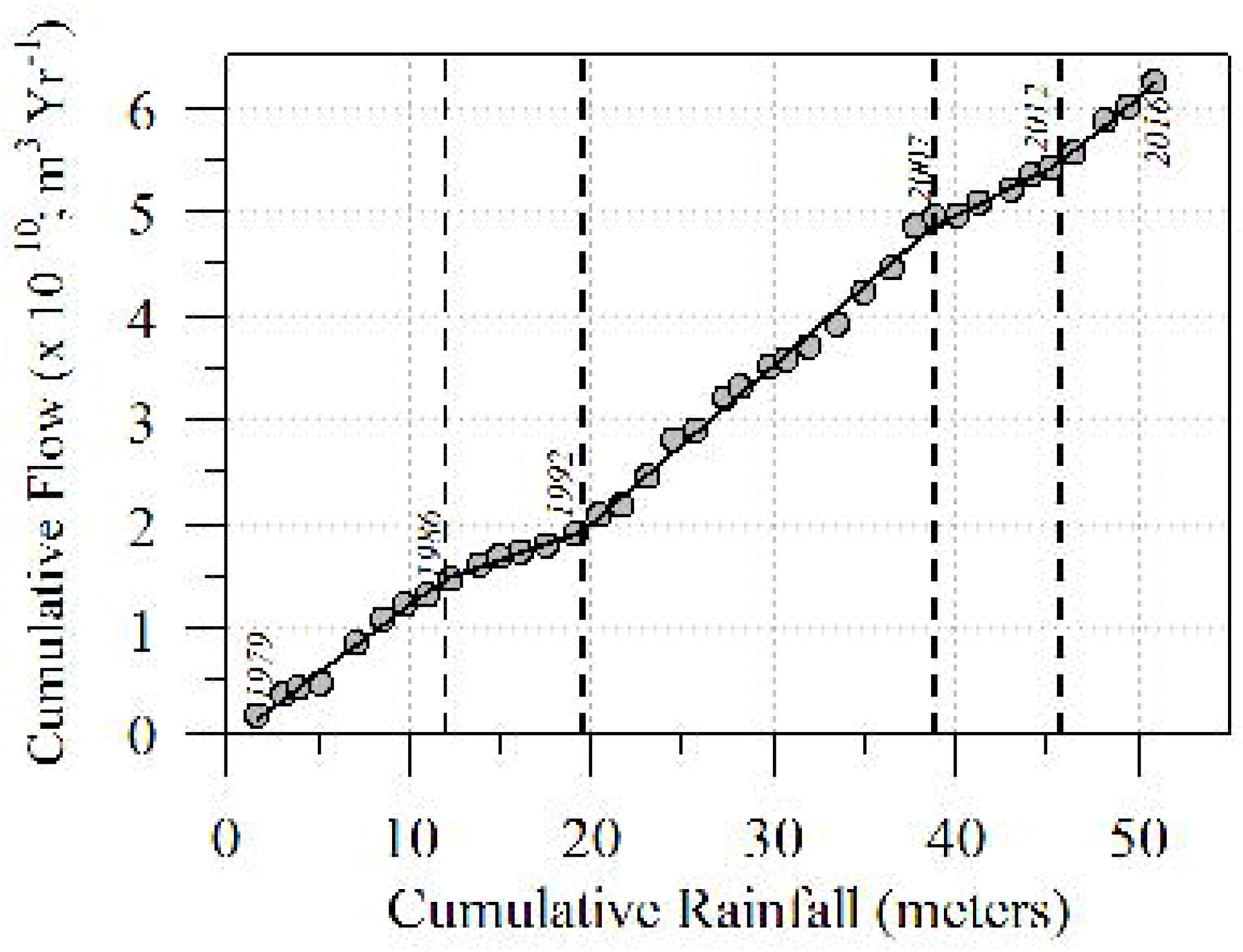
C-43 basin average cumulative rainfall versus S-79 cumulative flow from WY1979 to WY2016 (May 1, 1978 - April 30, 2016). Piecewise regression line is depicted by the black line through the points and breakpoints are identified by the vertical dashed lines. These breakpoints correspond to changes in water management practices and extreme climatic events (i.e. changes in rainfall).

### C-43 Canal-Caloosahatchee River Estuary Water Quality

Annual nutrient loads from Lake Okeechobee to the C-43 basin via the S-77 structure range from 4,925 to 361,792 kg yr^-1^ TP and 86,437 to 4,307278 kg yr^-1^ TN across POR (Fig. 4 and 5) with a mean of 63,699 ± 10,884 kg yr^-1^ TP and 1,231,271 ± 176,718 kg yr^-1^ TN, respectively. Annual HLR was significantly positively correlated with annual TN (r = 0.98, ρ<0.001) and TP (r = 0.95, ρ<0.001) loads at the S-77 structure. Meanwhile, annual nutrient loads at the S-79 range from 24,473 to 535,958 kg yr^-1^ TP and 138,759 to 6,532,271 kg yr^-1^ TN throughout the POR (Fig. 4 and 5). The mean POR nutrient loads at the S-79 were 224,571 ± 18,906 kg yr^-1^ TP and 2,588,347 ± 235,791 kg yr^-1^ TN, respectively. Period of record mean TP load from the S-79 was approximately 4 times higher than POR mean TP load observed at S-77 while mean TN load at S-79 was approximately two-times greater than mean TN load at S-77. Annual HLR was significantly positively correlated with S-79 annual TN (r = 0.92, ρ<0.001) and TP (r = 0.89, ρ<0.001) loads. The correlation coefficients for annual TN and TP loads versus HLR were lower for the S-79 than the S-77 suggesting more variability in the load and HLR relationship at the S-79.

**Figure 4.**
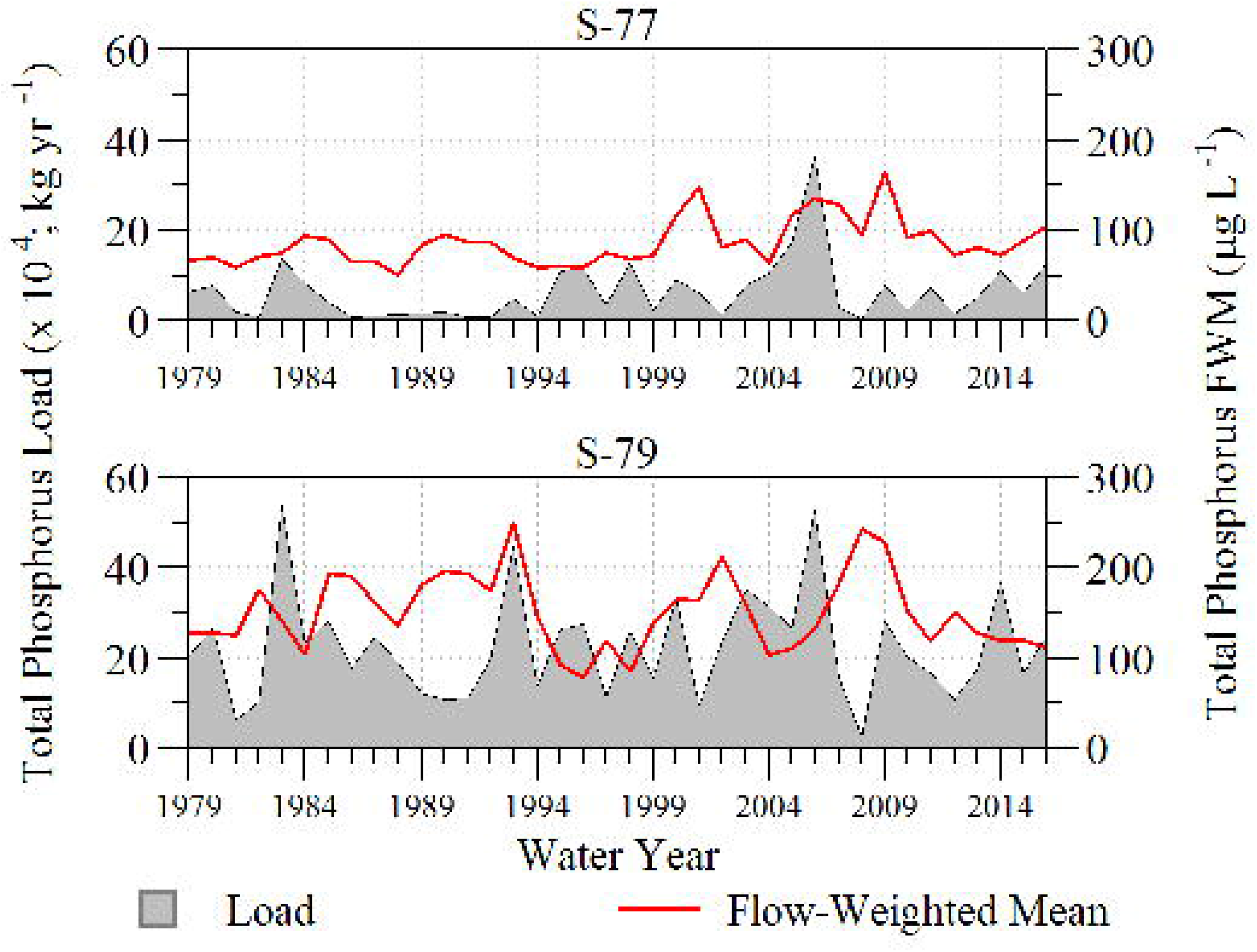
Annual total phosphorus load and annual flow-weighted mean for the S-77 and S-79 structures of the C-43 canal and Caloosahatchee river estuary between WY1979 and WY2016 (May 1, 1978 – April 30, 2016).

At the S-77 structure, annual FWM TP concentrations ranged from 51 to 165 μg L^-1^ with a mean of 86 ± 4.2 μg L^-1^ across the POR exhibiting a statistically significant increase (τ=0.31, ρ<0.001). A breakpoint was observed in FWM TP concentrations at WY1999 (K=249, ρ<0.01) concurrent with a significant change in flow volume timing through the S-77 (Fig. 4). This breakpoint corresponds to a period of high tropical activity followed by drought and eventual changes in water management regulation schedules for Lake Okeechobee. Furthermore, monthly TP FWM concentrations exhibited a seasonal pattern throughout the POR with higher FWM concentrations occurring in the late “wet” season (August and September) and lower concentrations during the “dry” season months between November to April (Supplemental Fig. 4). Annual TN FWM concentrations significantly decreased over the POR (τ=-0.27, ρ<0.05) ranging from 1.27 to 2.79 mg L^-1^ (Fig 5) with a POR mean of 1.71 ± 0.06 mg L^-1^. No statistically significant breakpoint was detected (K=170, ρ=0.09) in annual TN FWM concentrations at the S-77 structure.

**Figure 5.**
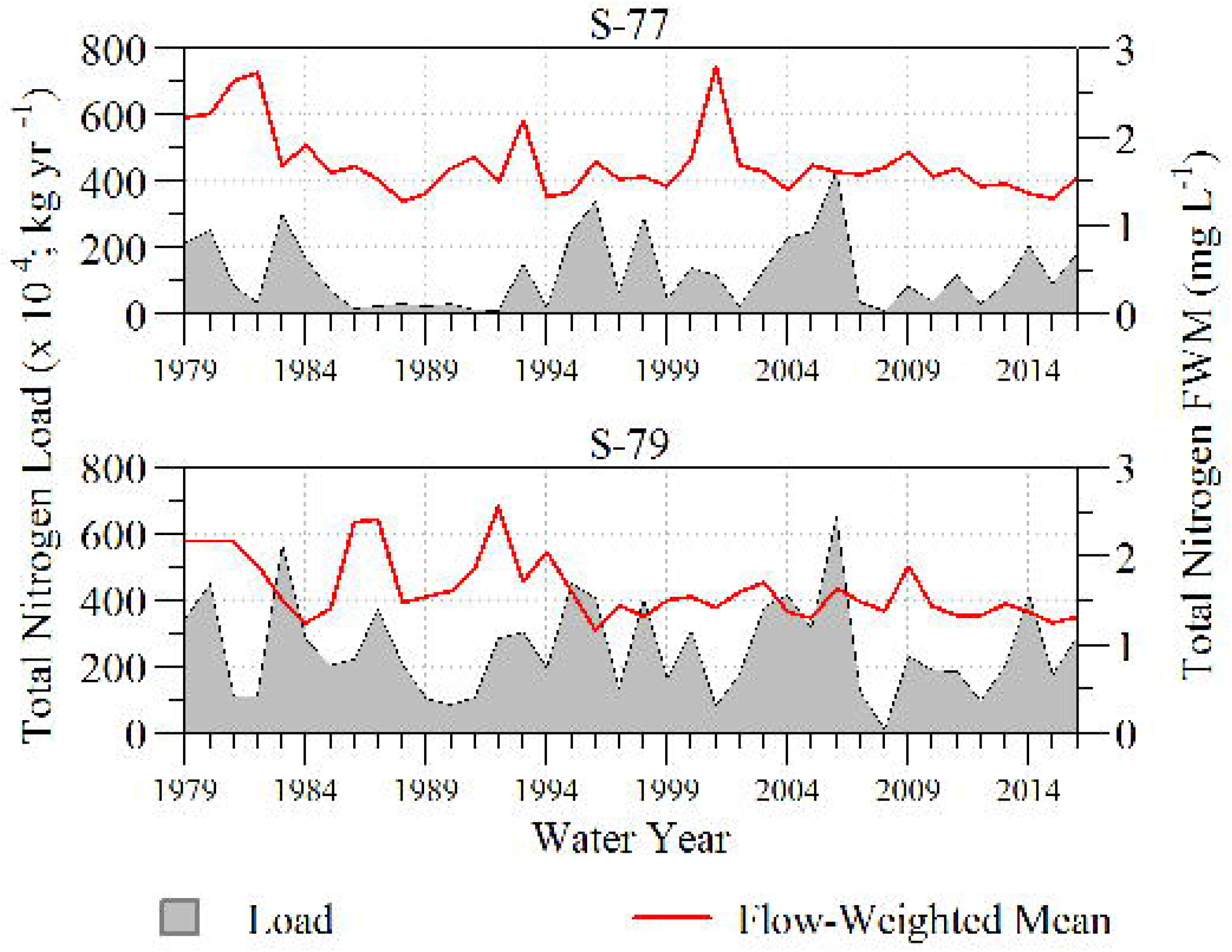
Annual total nitrogen load and annual flow-weighted mean for the S-77 and S-79 structures of the C-43 canal and Caloosahatchee river estuary between WY1979 and WY2016 (May 1, 1978 – April 30, 2016).

Further downstream at the S-79, annual TP FWM concentrations ranged from 78 to 250 μg L^-1^ with a mean of 151 ± 6.9 μg L^-1^ for the 37-year period. However, due to inter annual variability annual FWM concentration did not exhibit a statistically significant trend through time (τ=-0.08, ρ=0.47) (Fig. 4) and no significant breakpoints detected (K=126, ρ=0.37). Similar to the S-77, the S-79 exhibited a seasonal fluctuation in TP FWM concentrations during the period record with concentrations peaking during the June, July and August timeframe (Supplemental Fig. 4). Annual TN FWM concentrations ranged from 1.17 to 2.57 mg L^-1^ with a mean of 1.64 ± 0.06 mg L^-1^ during the POR. In contrast to TP, TN exhibited a statistically significant declining trend (τ=-0.39, ρ<0.001) at the S-79 structure (Fig. 5). A statistically significant breakpoint was detected for annual TN FWM concentration after WY1995 (K=245, ρ<0.01) where between year variability decreased resulting in more consistent concentrations with less between year variability (Fig. 5). Total Nitrogen FWM concentrations for the S-79 were similar to that of the S-77 with a couple of anomalous WYs that deviate from POR data (Supplemental Fig. 4).

### Nutrient Stoichiometry and Water Quality Longitudinal Analysis

The longitudinal water quality gradient along the CRE was evaluated in two ways, the first compared mean differences between freshwater, estuarine and marine regions over the entire POR and the second compared the steepness of the gradient on an annual basis. Along the C-43 Canal and CRE, water quality differed between regions (i.e. freshwater, estuary and marine). Nitrogen (i.e. TN and NH_4_), phosphorus (TP and OPO_4_) and TOC significantly declined along the freshwater to marine gradient (Table 2) between WY1999 and WY2016. Orthophosphate followed the same trend as TP, therefore the remaining analysis will focus on TP. Stoichiometric ratios of TN:TP ad OC:TP were significantly different between all regions of the estuary (Table 2), while OC:N ratios were not statistically different between regions (Table 2).

**Table 2.** Mean, Standard Error (Mean ± SE) and statistical comparison of water quality parameter for each region of the Caloosahatchee River estuary for data collected between May 1, 1999 and April 30, 2016. Values with different letters indicate statistically significant difference according to Dunn’s Multiple Comparison test using a critical value of α=0.05. Nutrient ratios (N:P, OC:N, OC:P, DIN:DIP, OC:DIP and OC:DIN) are represented as molar ratios.

| Parameter | Units | Freshwater | Estuary | Marine | Kruskal-Wallis Statistics |  |  |
| --- | --- | --- | --- | --- | --- | --- | --- |
| | | | | | $\chi^2$ | df | p-value |
| Total Nitrogen | mg L<sup>-1</sup> | 1.29 ± 0.02 (A) | 0.85 ± 0.03 (B) | 0.44 ± 0.02 (C) | 91.86 | 2 | <0.01 |
| Ammonia | mg L<sup>-1</sup> | 0.073 ± 0.003 (A) | 0.058 ± 0.002 (B) | 0.051 ± 0.001 (C) | 57.28 | 2 | <0.01 |
| Nitrate-Nitrite | mg L<sup>-1</sup> | 0.19 ± 0.012 (A) | 0.08 ± 0.005 (B) | 0.02 ± 0.002 (C) | 92.95 | 2 | <0.01 |
| Total Phosphorus | μg L<sup>-1</sup> | 133.3 ± 4.5 (A) | 96.1 ± 3.7 (B) | 52.5 ± 0.5 (C) | 80.54 | 2 | <0.01 |
| Ortho Phosphate | μg L<sup>-1</sup> | 89.2 ± 3.7 (A) | 60.0 ± 2.0 (B) | 40.1 ± 0.1 (C) | 82.83 | 2 | <0.01 |
| Total Organic Carbon | mg L<sup>-1</sup> | 16.69 ± 0.21 (A) | 11.89 ± 0.82 (B) | 7.25 ± 2.61 (C) | 83.58 | 2 | <0.01 |
| N:P | unitless | 26.99 ± 0.41 (A) | 20.57 ± 0.69 (B) | 13.56 ± 0.69 (C) | 21.69 | 2 | <0.01 |
| OC:P | unitless | 29.35 ± 0.53 (A) | 30.52 ± 4.43 (B) | 33.24 ± 12.76 (C) | 21.452 | 2 | <0.01 |
| OC:N | unitless | 1.08 ± 0.02 (A) | 2.01 ± 0.50 (A) | 2.92 ± 1.29 (A) | 0.35 | 2 | 0.84 |
| DIN:DIP | unitless | 8.96 ± 0.44 (A) | 7.69 ± 0.19 (B) | 7.48 ± 0.11 (B) | 9.82 | 2 | <0.05 |
| OC:DIP | unitless | 132.3 ± 1.3 (A) | 131.0 ± 17.5 (B) | 130.5 ± 48.7 (C) | 24.30 | 2 | <0.01 |
| OC:DIN | unitless | 15.78 ± 0.44 (A) | 16.59 ± 2.04 (A) | 17.94 ± 6.82 (B) | 16.70 | 2 | <0.01 |

The annual mean TN:TP differed along the CRE with freshwater portions exhibiting a clear spatial trend with freshwater portion above a molar ratio of 20:1 (Guildford and Hecky 2000) similar to the Redfield ratio (16:1) derived for dissolved N:P in oceanic systems (Redfield 1958), estuarine portions were approximately 20 while marine were below 20 (Table 2). This suggests that the CRE transitions from a potentially P-limited system in the freshwater portion to potential N-limitation in the marine portion, with the estuarine shifting between N and P limiting depending on the influence of tidal and freshwater flow. The variability in annual mean TN:TP were greater in marine (variance: 13.5; range: 8.0– 19.0) and estuarine (variance: 45.2; range: 7.7 – 35.0) portions of the CRE with marine portions always being below a TN:TP ratio of 20 and in some years the estuarine portion experiencing TN:TP ratios less than 20. Meanwhile freshwater (variance: 7.5; range: 24.6 – 32.7) portions of the CRE were well above the TN:TP ratio of 20 (Table 2).

Trends in inorganic nutrient stoichiometry differed slightly from bulk nutrient stoichiometry. Most notably annual mean DIN:DIP ratios were below the Redfield 16:1 ratio (Redfield 1958) and slightly above the 5:1 ratio suggested by (Doering et al. 2006). However there are instances when DIN:DIP ratios fell below the 5:1 DIN:DIP ratio in freshwater and estuarine portions of the CRE (Supplemental Fig. 5). Annual mean DIN:DIP ratios were statistically similar between estuary and marine portions of the CRE and statistically different from that of the freshwater portion. Annual mean OC:DIP molar ratio differed significantly along the CRE and OC:DIN ratios were statistically similar between freshwater and estuarine portions but statistically different from the marine portion.

Along the gradient of fresh to marine waters TP, TN and TOC decreased (Fig 6). A total of 12 years of annual mean TP data were used in the longitudinal gradient analysis, however, WYs 2003 to 2006 and 2008 were excluded due to the lack of suitable number of samples and representative stations. Annual mean TP concentrations are highly variable in the freshwater and estuary reaches while concentrations observed at marine stations are consistent from year to year (Fig. 6). Concentration gradient rates (i.e. slopes) range from -3.4 to -1.1 μg L^-1^ TP km^-1^ with rates generally increasing over time (Fig. 6). However, due to the high variability (variance: 0.46 μg L^-1^ TP km^-1^) the trend in concentration gradient rates was not statistically significant (τ=0.12, ρ=0.63). Similar to the TP longitudinal gradient analysis, 12 years of annual mean TN data were used. Concentration gradient slopes ranged from -0.03 to -0.02 mg L^-1^ TN km^-1^ with slopes increasing over time (τ=0.55, ρ<0.05; Fig. 6). A total of 11 years of annual mean TOC data was used in the longitudinal gradient analysis specific to TOC, WYs 2003 to 2008 were excluded from the analysis due to lack of a suitable number of samples and representative stations. Concentration gradient rates range from -3.8 to -2.4 mg L^-1^ TOC km^-1^ with no significant temporal trend (τ=-0.31, ρ=0.21; Fig. 6). Furthermore, TP (r=0.81, ρ<0.001), TN (r=0.84, ρ<0.001) and TOC (r=0.85, ρ<0.001) were positively correlated with monthly mean CDOM at the RECON-water quality paired sites along the CRE.

**Figure 6.**
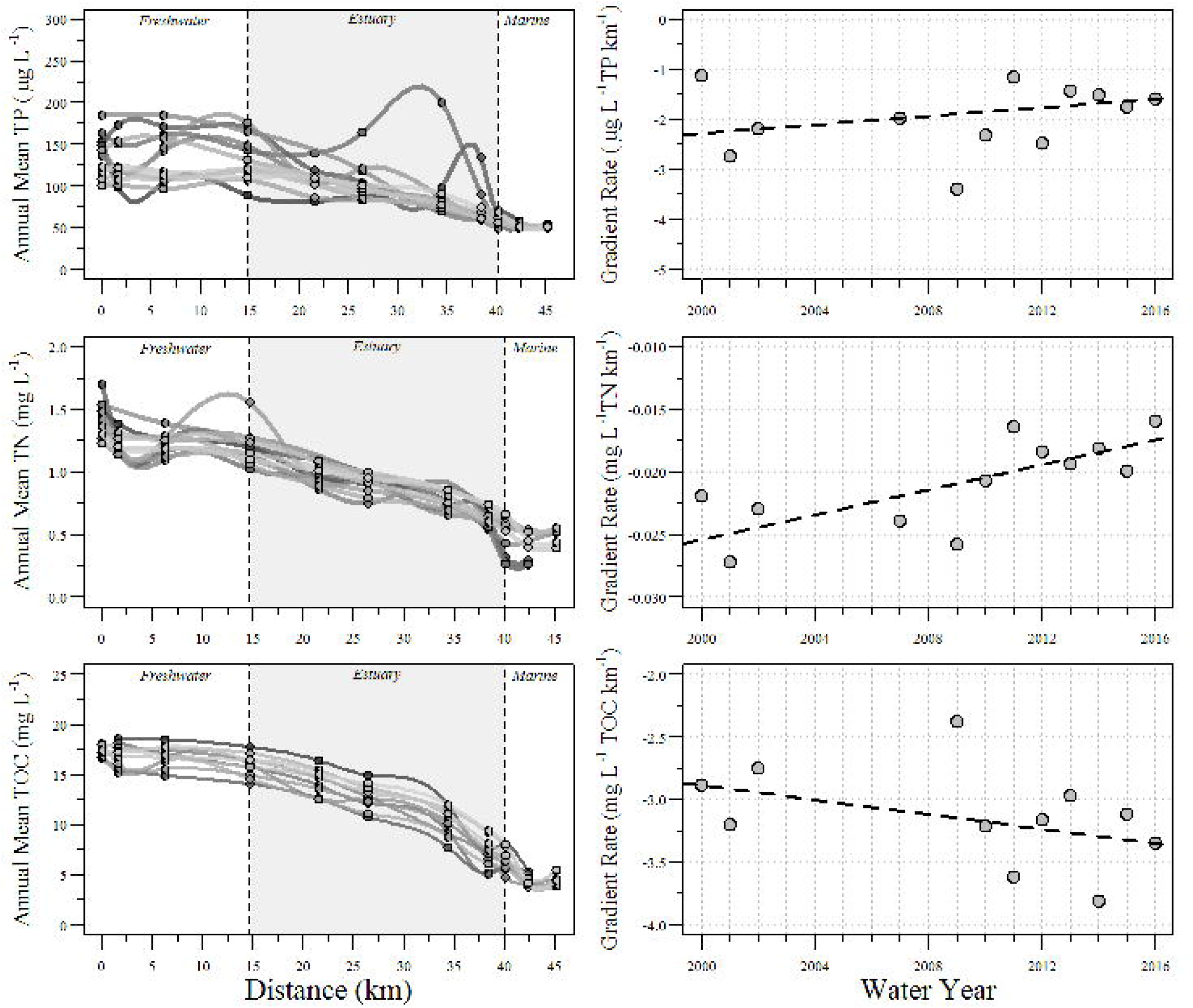
Left: Annual mean Total Phosphorus (TP), Total Nitrogen (TN) and Total Organic Carbon (TOC) along the Caloosahatchee River Estuary (CRE) between water year (WY) 2000 and 2016 (May 1, 1999 – April 30, 2016). Distance zero is the S-79 structure, the start of the freshwater portion of the CRE. Dark lines indicate earlier water years while light lines indicate later years. Right: Concentration gradient rate for each parameter by WY between WY2000 and 2016, dashed black line indicates Kendall trend using the Thiel-Sen estimate.

### Aquatic Metabolism and Influencing Factors

Between WY2009 and WY2016 the Ft Myers, Shell Point and GOM monitoring locations had adequate data to calculate aquatic metabolism for all WYs during this period. However, data for Beautiful Island was limited to WY2013-2016 and Tarpon Bay was limited to WY2011-2016. During the POR daily NAP estimates ranged from -8.4 to 8.2 g O_2_ m^-2^ d^-1^, GPP estimates ranged from -18.8 to 14.0 g O_2_ m^-2^ d^-1^ and ER estimates ranged from -16.9 to 20.3 g O_2_ m^-2^ d^-1^ for the five RECON sites used in this study.

The CRE is predominately heterotrophic with respect to aquatic metabolism in that all sites within the CRE had a statistically significant GPP:ER ratio less than one indicating that GPP < ER (Table 3). Daily GPP was significantly different between marine and estuarine regions (χ^2^=304.32, df=1,ρ<0.001), with GPP in marine (1.90 ± 0.03 g O_2_ m^-2^ d^-1^) portions being greater than estuarine (1.41 ± 0.02 g O_2_ m^-2^ d^-1^) portions of the CRE. Meanwhile, daily ER estimates were significantly greater in estuarine portions (-1.64 ± 0.02 g O_2_ m^-2^ d^-1^) relative to marine portions (-1.96 ± 0.02 g O_2_ m^-2^ d^-1^) of the CRE (χ^2^=138.61, df=1,ρ<0.001). As a result of the balance between GPP and ER, daily NAP estimates were significantly greater in marine portions (-0.07 ± 0.01 g O_2_ m^-2^ d^-1^) relative to estuarine portions (-0.24 ± 0.01 g O_2_ m^-2^ d^-1^) of the CRE (χ^2^=341.41, df=1,ρ<0.001).

**Table 3.**
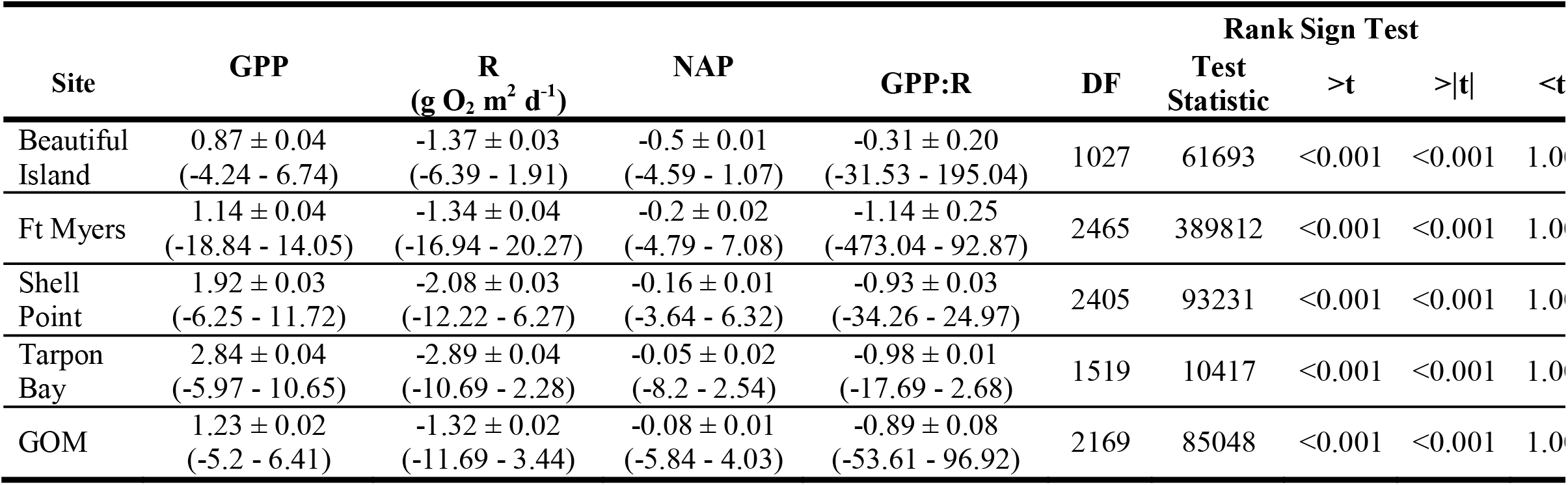
Mean ± standard error (minimum – maximum) gross primary productivity (GPP), respiration (R), net aquatic productivity (NAP) and GPP: R ratio estimates. Wilcoxon rank sign nonparametric test statistic results comparing site daily GPP:R values to a value of zero (μ=0).

Daily NAP was negatively correlated with TP (r=-0.31, ρ<0.001), TN (r=-0.41, ρ<0.001), TOC (r=-0.37, ρ<0.001), TN:TP molar ratio (r=-0.40, ρ<0.001) and TOC:TP molar ratio (r=-0.28, ρ<0.01) but not correlated with TOC:TN molar ratio (r=-0.03, ρ=0.76). Change-point analysis identified statistically significant breakpoints for TP, TN and TOC when compared to NAP. A statistically significant breakpoint was identified in the relationship between NAP and TP (R^2^ =0.08, F_(3,100)_=2.71, ρ<0.05) occurring at 53 μg L^-1^ which corresponds with the P boundary between marine and estuary suggesting different controls of net productivity between regions (Fig. 7). Additionally, a statistically significant breakpoint was identified in the relationship between NAP and TN (R^2^ =0.12, F_(3,100)_=4.54, ρ<0.01), however the breakpoint identified an area where only two other samples occurred after the breakpoint at a TN concentration of 0.65 mg L^-1^ (Fig. 7). Even though the piecewise regression indicated a significant change in slope, the R^2^ was relatively low and the breakpoint occurred in an area of decreased data density to the point that the line after the breakpoint would be unrepresentative. Therefore, the NAP-TN breakpoint should be viewed with caution, additional investigation is needed to further elucidate the interaction between TN and NAP in marine and estuarine systems such as the CRE. A statistically significant breakpoint was identified in the relationship between NAP and TOC (R^2^ =0.11, F_(3,100)_=4.06, ρ<0.01) at a TOC concentration of 11.89 mg L^-1^ (Fig. 7).

**Figure 7.**
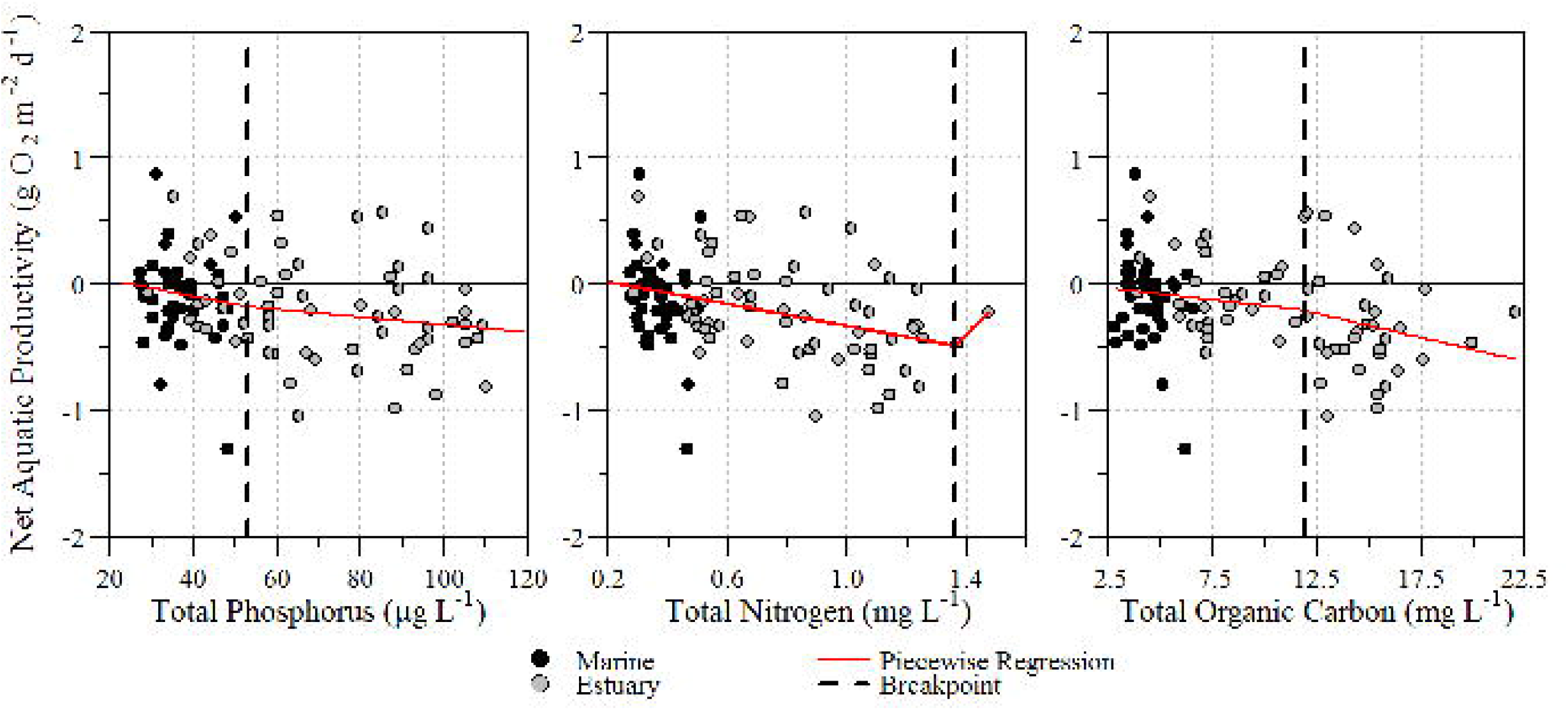
Piecewise regression of water quality parameters and net aquatic productivity along the Caloosahatchee River Estuary (CRE) between water year (WY) 2008 and 2016 (May 1, 2007 – April 30, 2016) using five River, Estuary and Coastal Observing Network (RECON) sites and associated paired ambient water quality stations. Piecewise regression lines and associated breakpoints are identified in soil red line and vertical dashed black line, respectively.

## Discussion

### Hydrologic dynamics within C-43 Canal-Caloosahatchee River Estuary aquatic continuum

Overall the C-43 Basin and the CRE have undergone numerous alterations to facilitate navigation, flood control and regulatory releases from Lake Okeechobee (Qiu and Wan 2013). The conglomeration of these modification in combination with climatic shifts and changes in upstream water management may confound the interpretation of statistically significant changes observed in flow dynamics at structures along the C-43 canal and into the CRE. The observed change in monthly flow volume from the S-77 during WY1994 marks a period where one of the first Lake Okeechobee regulation schedules known as Operational Schedule 25 or “Run 25” became effective (Fig. 8). The “Run 25” regulation schedule represented a trip-line type of operation where if the lake water elevation passes a regulation line, action was taken. During “Run 25” water levels in Lake Okeechobee were managed from regulatory (flood-control) and non-regulatory (primarily water supply) purposes while attempting to maintain a relatively high stage for prolonged periods of time. This high water level within the lake resulted in unintended impacts to lake ecology (SFWMD 2005; SFWMD and USACE 1999; Vearil 2008). The intra- and inter-annual variability observed in the flow volumes (Supplemental Fig. 2 and Fig. 2) suggests a high degree of variability reflective in water management associated with water needs in either the upstream or downstream waterbodies rather than seasonal hydrologic management. The pattern and magnitude of flow observed at the S-77 is influenced by the subtropical climate of south Florida regulatory discharges from Lake Okeechobee and withdrawals for irrigation and water supply for agricultural and urban uses (Qiu and Wan 2013).

**Figure 8.**
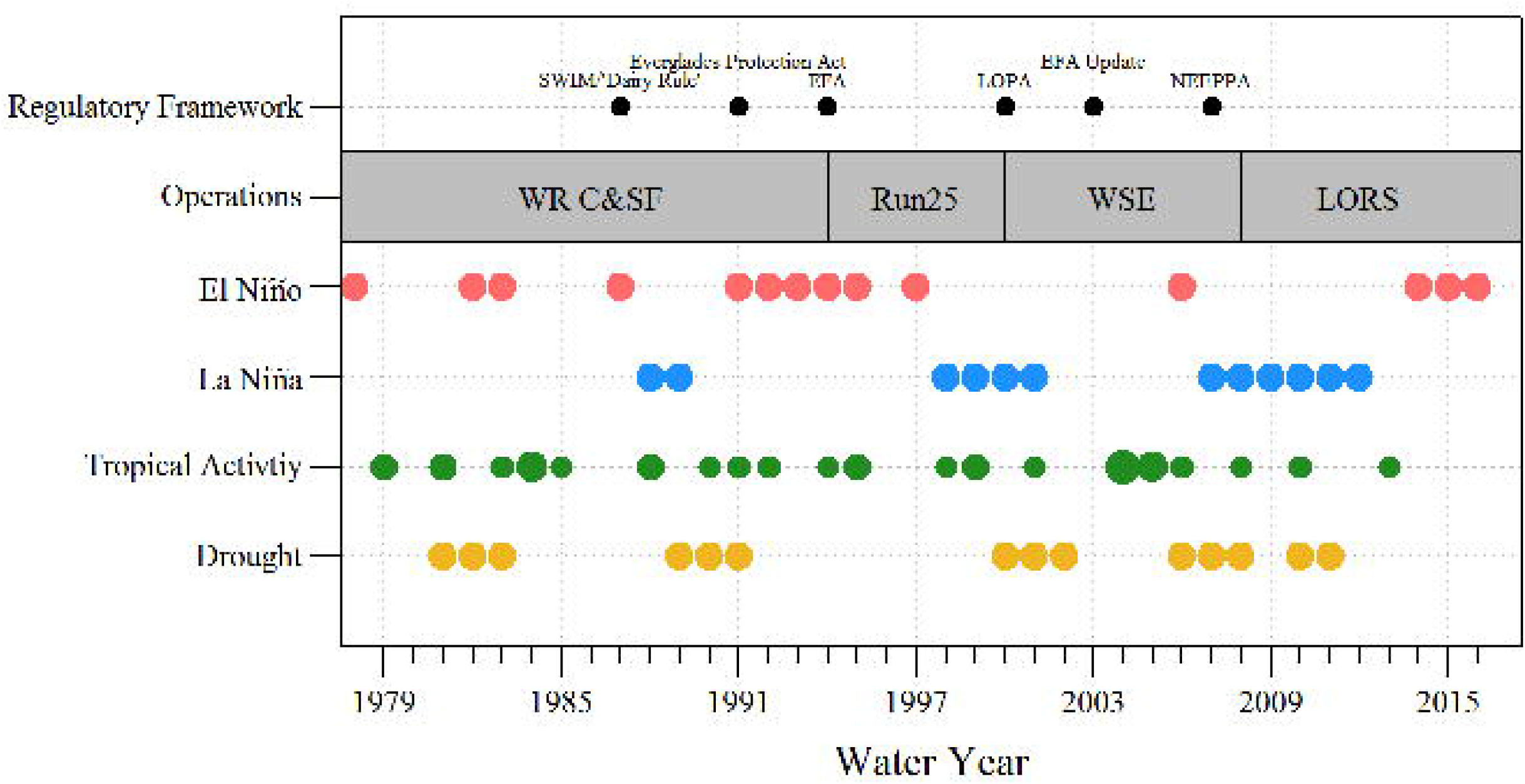
Lake Okeechobee operation timeline relative to climatic events and regional regulatory action. El Niño (red point) and La Niña (blue points) occurrence (Data Source: <u>http://www.cpc.noaa.gov</u>), Tropical activity with relative size of the point indicating the number of storms (Data Source: <u>https://coast.noaa.gov</u>) and identification of drought years (yellow points; Data Source: (Abtew et al. 2009; Abtew and Ciuca 2017). SWIM=Surface Water Improvement and Management; ‘Dairy Rule’ authorized by 62-670.500 Florida Administrative Code; EFA=Everglades Forever Act (Everglades Protection Act and EFA authorized by § 373.4592 Florida Statute); LOPA=Lake Okeechobee Protection Act NEEPPA=Northern Everglades and Estuaries Protection Program (LOPA and NEEPPA authorized under § 373.4595 Florida Statute).

Estimated annual total flow from the C-43 basin was greater than flow from Lake Okeechobee (i.e. S-77) indicating that on average, basin flow rather than lake management contributed to excessive freshwater flows to the estuary. Furthermore, the double-mass curve analysis (Fig. 3) identified several changes in the discharge run-off relationship within the C-43 basin to the estuary via the S-79, some of these changes corresponded with upstream management adjustments while other resulted from changes in climate (Fig. 8, Supplement Fig. 3). The period of 1992 to 2007 included El Nino weather patterns and several hurricanes and tropical storms making landfall or passed near southwest Florida (Fig. 8), resulting in significant contributions of rainfall. Due to the large size of the C-43 basin (watershed) and modification to the watershed to facilitate drainage and irrigation, rainfall-driven watershed run-off is a significant contributor of flow volume to the estuary. Additionally, during this period several upstream operational changes came into effect including for Lake Okeechobee implementation of “Run 25” mentioned earlier, the Water Supply and Environmental (WSE) Operational Schedule which came into effect in 2000 and the implementation of the Lake Okeechobee Regulation Schedule (LORS) effective 2008 (SFWMD and USACE 1999; Vearil 2008) (Fig. 8). Prior to May 2000, the Lake experienced prolonged water depths. Therefore, in an attempt to restore ecological function to the Lake, a managed recession of occurred in April-May 2000 followed by a prolonged drought (Fig. 8) therefore WSE was not officially implemented until after the drought broke in 2003. Similarly, during 2006-2008 timeframe, a severe drought prolonged the official implementation of LORS and was not initiated until after 2008 (Fig. 8)(Abtew et al. 2009).

Freshwater flow to the CRE is a combination of several upstream dynamics. These dynamics are driven in part by long- and short-term climate which affects water management for ecological, flood protection and water supply purposes. The operation of Lake Okeechobee and flows from the C-43 basin are at times decoupled due to regional climate and rainfall and upstream water management.

### Freshwater Aquatic Environment and Watershed Dynamics

Generally, nutrient loads were greater at S-79 than at S-77 suggesting that the C-43 basin, at times can act as a nutrient source however it is uncertain if this is due to legacy nutrients in the river and canal bottom, high nutrient ground water upwelling, basin sourced nutrient run-off or a mixture of these factors. Furthermore, both the S-77 and S-79 TP and TN loads exhibited large inter-annual variations predominately driven by changes in flow volumes in response to either changes in management strategies, climatic events or a combination of these factors (Fig. 4, 5 and 8). As seen in larger river systems flow and watershed run-off drive trends in water quality with respect to loads (i.e. flow and concentration) (Hagy et al. 2004; Perona et al. 1999; Shrestha and Kazama 2007; Turner and Rabalais 2003). The CRE upstream watershed is predominately agricultural lands (~40%) which both creates a significant demand for water during the dry season and becomes a source of stormwater run-off and nutrients during the wet season influencing the water quality inputs to the CRE (Qiu and Wan 2013).

In the Chesapeake Bay, year-to-year variability of flow and associated nitrate load from the Susquehanna River in addition to watershed level effects accounted for changes in downstream biological response (i.e. volume of hypoxic water; (Hagy et al. 2004). Turner and Rabalais (2003) have suggested that conversion of lands to intense farming in the Mississippi river watershed over the past two-centuries has significantly altered water quality downstream. In the warm and dry Mediterranean climate, seasonal increases in human activates such as domestic water use and agricultural activates have been linked to high nutrient concentrations in the Alberche River located in Spain (Perona et al. 1999). In the temperate Fuji River watershed of Japan, variation in water quality was related to flow, water temperature, and nutrient inputs from different regions of surrounding watersheds with a variety of land uses (Shrestha and Kazama 2007). Across these systems, flow volume is the primary influence on trends in water quality downstream.

Change points in annual TP and TN FWM concentrations at both the S-77 and S-79 (Fig. 4 and 5) occurred during or after periods of changes in upstream operations (see previous discussion), climatic events, changes in off-site agricultural management practices such as best management practices (BMPs) or implementation of regulatory framework (Fig. 8). The differences in water quality change points is partly reflective of changes in Lake regulation schedules, climatic events and regional climate patterns resulting in variability in runoff volumes from the C-43 basin altering the timing of TP delivered to the C-43 canal and subsequent CRE. However, the change in TN FWM concentrations at the S-77 seemed to be associated with changes in climatic conditions within the region and implementation of several environmental regulatory frameworks used to address water quality issues or a combination of both. These regulatory frameworks include the Florida Surface Water Improvement and Management (SWIM) act of 1987 which required the implementation of agricultural BMPs. In addition to the SWIM act of 1987, the state of Florida promulgated the “Dairy Rule” in 1987 (62-670.500 Florida Administrative Code) which reduced run-off from dairy operation entering waterways by requiring waste water treatment systems in an attempt to reduce nutrient loads (Anderson and Flaig 1995). In addition to SWIM and the Dairy Rule other regulatory frameworks were authorized by the State of Florida focusing on the development, implementation and regulation of BMPs including the Everglades Protection Act, Everglades Forever Act, Lake Okeechobee Protection Act and the Northern Everglades and Estuaries Protection Program (Fig. 8).

The variability of TP FWM concentration at the S-79 structure could be linked to the “flashiness” of the C-43 basin facilitated by watershed modifications to promote drainage and water control (Cook 2014). The CRE and C-43 canal were linked to Lake Okeechobee through the evolution of the Central and South Florida Flood Control Project (C&SF) during the late 1800’s and early 1900’s with subsequent modifications to improve navigation and recreation in the mid 1900’s. The features of the C&SF project include a system of canals, water control structures and storage areas spanning the entirety of southern and central Florida. In addition to changes of hydrology from the C&SF project a multitude of other structural and physical alterations has occurred in the CRE and C-43 basin. These alterations resulted in changes to the historic hydrologic conditions of the watershed due to secondary and tertiary canals that connect to both the C-43 canal and CRE. These canals provide navigational access and convey water for both drainage and irrigation purposed to accommodate agricultural and urban areas (Buzzelli et al. 2016). Due to these canals, in combination with increased development and impervious surfaces stormwater runoff volumes have increased relative to predevelopment conditions. Therefore, the variability of TP FWM concentrations at the S-79 structure could be due to several factors including contributions of flow dependent changes in P concentrations due to differences in volumes and sources of water (i.e. stormwater run-off, upstream lake management, etc.).

In addition to the “flashiness” of the basin the seasonality of TP FWM concentrations are somewhat out of sync of what one would intuitively think. During any given WY, the peak monthly TP FWM concentrations occur first at downstream water control feature (i.e. S-79) early in the wet season (i.e. May – August) presumably in response to run-off from the upstream basin. Meanwhile, peak monthly TP FWM concentrations at the upstream water control feature (i.e. S-77) occur toward the end of the wet season (i.e. August – October). Timing of nutrient loads from the structures are similar to FWM concentration trends. This timing in nutrient delivery is likely important to the ecology of the river and downstream estuary with an immediate wet season pulse of nutrients at the onset of the wet season followed by a later (less intense) delivery of nutrients delivered from further upstream subsidizing nutrients required and are facilitated by retention time of the river (i.e. weeks to months), localized climatic events and water management.

### Nutrient dynamics along the aquatic continuum

Generally, nutrient limitation shifts along the reach of an estuary (R. Howarth et al. 2011). As waters transition from freshwater to brackish and finally to marine, the water quality characteristics also change due to sources of inputs (i.e. runoff), internal processes and outputs from the given system. Generally, the supply of P regulates the production in freshwater systems while N regulates production in estuarine and marine systems (Conley et al. 2009). Integral to the concept of nutrient limitation is the supply of one resource relative to another (i.e. stoichiometry), The difference in limiting nutrients between ecosystems could be linked to the availability of N, P or both (Doering et al. 1995). The identifiable stoichiometric relationship, known as the Redfield ratio, suggests that marine planktonic nutrient composition has a characteristic molar ratio and the abundance of nutrients are regulated by reciprocal interactions between organisms and the environment (Redfield 1958). The Redfield dissolved N: dissolved P molar ratio of 16:1 is often used as a benchmark for differentiating N from P growth limiting conditions for phytoplankton in marine environments (Geider and La Roche 2002; Redfield 1958). However when considering total nutrient values, a ratio of 20:1 has been used to indicate algal nutrient limitations (Guildford and Hecky 2000) and a ratio of 8:1 for microbial community nutrient limitation (Heil et al. 2007; McPherson and Miller 1990). It has also been suggested that total fractions rather than dissolved fractions are better indicators of trophic state and nutrient limitation as demonstrated in stream ecosystems (Walter K. Dodds 2003).

The aquatic continuum along the C-43 Canal/CRE followed the typical “right side-up” estuarine pattern (Childers et al. 2006b) with decreasing nutrient concentrations along the flow path with relatively higher nutrient concentrations at the estuary’s headwaters (i.e. freshwater reaches) and lower nutrient concertation along the downstream segments of the flow path. It is broadly assumed that N is the primary limiting nutrient in the marine portions of the CRE and P is the limiting nutrient in the freshwater (i.e. headwaters of the CRE and upstream of the S-79) portions of the CRE. The TN:TP molar ratio of 20 to indicate N-limitation suggests that freshwater and estuarine portions of the CRE are P growth limiting or N and P co-limiting. However, there are times in which the lower estuarine portions of the CRE becomes N-limited (i.e. TN:TP<20) consistent with results presented by (Doering et al. 2006). Furthermore, the TN:TP molar ratio of 8:1 suggests a potential P growth-limiting conditions to microbial communities in lower estuary and marine portions of the CRE (Supplemental Fig. 4). Near-shore at the terminus of the CRE, (Heil et al. 2007) observed low dissolved inorganic N:P (DIN:DIP) molar ratio of 4.5 and particulate N:P < 8 suggesting N growth limiting conditions for microbial communities. Low N:P molar ratios are not uncommon and have been observed in other ecosystems including Port Phillip Bay, Australia, other Australian rivers (i.e. Murray River, Victorian River) and the Peel-Harvey system in Washington (USA) with some studies suggesting trace metal availability rather than nutrient limitation regulates phytoplankton/algal blooms (Harris 2001) which could also be problematic to the CRE and near shore region. These potential nutrient growth limitations to the CRE is driven in large part by flow and nutrient regimes from upstream sources.

Freshwater discharge from rivers influences estuarine water quality through changes in salinity regimes, nutrient composition (organic versus inorganic), nutrient quantity (i.e. increased loads) and inputs of growth limiting nutrients altering the balance of nutrients potentially causing the occurrence of algal blooms, hypoxia, and other adverse effects to the ecosystem (W.K. Dodds 2006; Doering and Chamberlain 1999; Perez et al. 2010). Indirectly, freshwater discharges also influence estuarine water quality based on the effects of discharges on estuarine hydrodynamics, more specifically residence time. The change to residence time and its effect to water quality is two-fold, with lower residence times particulate nutrients have less time/distance to settle to the river bottom and lower residence times do not allow for sufficient time for biological uptake (Doering and Chamberlain 1999; Eyre 1998; Harris 2001; Paerl et al. 1998). The most notable difference in the CRE water quality over time is the relative increase in the slope of the TN longitudinal gradient (Fig. 6) over time even though TN concentrations entering the estuary have significantly declined across the long-term POR. While concentrations are on the decline entering the estuary, the slightly increasing trend of monthly flow could negatively impact the water retention time of the estuary, with concurrent point source increases TN load, and/or increased loads from run-off to the tidal basins (not accounted for in this study)(Buzzelli et al. 2016) could contributing to the increase in the TN gradient of the CRE. Thereby the interplay between water quality inputs, hydrodynamic variability and estuary dynamics all influence the water quality condition of the CRE.

### Estuarine Metabolism

Estuaries are generally net heterotrophic with respect to aquatic productivity, resulting in negative NAP estimates (Caffrey 2004; Caffrey et al. 2013). Caffrey (2004) estimated aquatic productivity across 22 National Estuary Research Reserves spanning five geographic regions using data from 42 monitoring sites and concluded that all but three were heterotrophic. Hoellein et al. (2013) also reported the majority of estuaries reviewed in their study were net heterotrophic. Much like other estuaries, the CRE is predominately heterotrophic with respect to aquatic productivity in that negative NAP estimates were observed at all sites within the CRE (Table 3). Aquatic productivity estimates observed in the CRE during this study were consistent with values observed in other estuarine and river systems as reported in previous studies (Caffrey 2004; Caffrey et al. 2013; Hoellein et al. 2013; Shen et al. 2015; J. Thébault et al. 2008).

Aquatic productivity can be limited by both nutrients and light (Petersen et al. 1997). Furthermore, a debate regarding which nutrient limits productivity and the roles that N and P play in limiting productivity across the aquatic continuum has been ongoing (R. Howarth and Paerl 2008; Paerl 2009; Schindler et al. 2008). Some studies have suggested that productivity can be limited by N or P but also co-limited by either (R. W. Howarth and Marino 2006; Lewis et al. 2011). In lake ecosystems TP and DOC concentrations were important drivers of aquatic productivity (Cole et al. 2000; del Giorgio and Peters 1994). Within the CRE the negative correlation of TP, TN and TOC could be the result of trade-offs between limiting factors of aquatic productivity. Both nutrients and highly colored freshwater could limit productivity in the CRE. However, more intensive analysis is needed to determine specific limiting factors related to light conditions, nutrient concentrations and freshwater influences on aquatic production in the CRE.

The relationship of NAP and TOC observed within the CRE is consistent with observations in lake ecosystems, where aquatic productivity changes occurred across OC concentration gradients. Hanson et al. (2003) observed a breakpoint in NAP estimates at a DOC concentration of approximately 10 mg L^-1^. Within the CRE, at low TOC concentrations GPP was relatively comparable to R which suggests that autochthonous carbon provides most of the liable substrate for aquatic respiration. Low TOC concentrations were generally associated with marine and tidally influenced estuary sites (i.e. Gulf of Mexico, Tarpon Bay and Shell Point) (Fig. 6) where frequent exchange of ocean water occurs and transformation of OC occurs due to changes in salinity (i.e. ionic strength). Meanwhile, high TOC concentrations and variable NAP were associated with more freshwater or brackish water sites (i.e. Fort Myers and Beautiful Island). Variability in the NAP and TOC relationship at these sites could be driven by OC from freshwater deliveries to the estuary, transformation of OC due to changes in ionic strength, infrequent tidal flushing high in the estuary and local tidal basin run-off. The difference between OC change points observed in lake (Hanson et al. 2003; Prairie et al. 2002) and estuary (this study) ecosystem could also be due to differences in hydrology and TP concentrations. The lake systems studied by previous researchers (Hanson et al. 2003; Prairie et al. 2002) observed TP concentrations <20 μg L^-1^, while TP concentrations within this study were >20 μg L^-1^ (Fig. 7). Organic C has been shown to increase heterotrophic respiration and microbial productivity, but OC can also limit productivity via shading of autotrophic organism via increased CDOM or self-shading via increased biomass (Hanson et al. 2003; Hoellein et al. 2013; Karlsson et al. 2009; Maranger et al. 2005). Therefore, a region-specific trade-off between CDOM, OC, TP and aquatic productivity across aquatic ecosystems within the aquatic continuum where water column parameters directly (nutrient limitation) or in-directly (shading due to high OC) regulate productivity could occur. Regardless more research is needed to further explore the role of nutrients and OC in regulating productivity across the aquatic continuum to better understand and protection flora and fauna in a high managed system.

### The Caloosahatchee River Restoration Effort

As part of the Comprehensive Everglades Restoration Plan (CERP) the CRE and surrounding estuaries were identified as regions of interest. Restoration of these systems to a healthy, productive aquatic ecosystem is important to maintaining the ecological integrity of the region (U.S. Army Corps of Engineers and South Florida Water Management District 2010). To restore the ecological integrity of the CRE, several alternative restoration plans were considered all emphasized storage of water upstream of the estuary during the wet season to prevent or reduce high discharge events and treatment of water to reduce nutrient loads to the estuary.

The selected project included a deep reservoir with 0.21 km^3^ (170,000 acre-feet) of storage. This storage capability will allow the reduction of high flow events and greater control of freshwater releases to the estuary and help improve water quality. Based on modeling efforts to support the selected restoration plan, weekly high flow characterized as volumes >127 m^3^ s^-1^ (4,500 ft^3^) could be reduced by approximately 80% and nutrient (i.e. TP and TN) and sediments loads to the estuary will be reduced by approximately 30% (National Research Council 2010). Furthermore, modeling suggests an overall improvement to the estuarine environment including estuarine indicator species such as oysters and seagrasses (U.S. Army Corps of Engineers and South Florida Water Management District 2010). Based on this information the implementation of the CERP project for the CRE/C-43 would result in the reduction of flow and nutrient load to the estuary which in turn could reduce the flow-weighted mean nutrient concentrations. Assuming that a freshwater condition still exists at the CRE headwater (i.e. S-79), the same longitudinal gradients observed during this study should persist however the rate of change in nutrient concentrations along the gradient should hypothetically decline due to the reduction of nutrients entering the estuary via the S-79, assuming the marine end-member of the gradient doesn’t change. This reduction in nutrients and volume could alter nutrient limitations within the CRE along the aquatic continuum and potentially reduce the variability observed in this study.

## Conclusions

Upstream water management and watershed characteristics are profound drivers of downstream estuarine ecology. With respect to the CRE, modifications to the C-43 basin have significantly altered the flow of water and nutrients through the basin and into the estuary resulting in large inputs of runoff from the terrestrial ecosystem upstream of the CRE (starting point of the aquatic continuum). Rainfall and run-off from the C-43 basin accounts for the majority of flow into the CRE rather than water management associated with Lake Okeechobee. Furthermore, the data presented in this study indicates that the C-43 basin contributes more nutrients to the CRE than Lake Okeechobee. However, changes in water quality conditions in the C-43 and CRE have changed through time, responding to changes in upstream water management and regulatory action intended to improve water quality conditions for the upstream waterbodies (i.e. Lake Okeechobee), C-43 basin and CRE. Upstream water management plays a significant role in regulating ecosystem level productivity due to changes in available TP, OC, fresh/saline water balance and dissolved organic matter.

This work helps to better understand the potential impact of upstream water management, basin level actions and local mechanisms on water quality and ecological integrity of the CRE by applying an aquatic continuum approach. Because this approach allows for independent analysis of each region (i.e. freshwater, estuarine, marine) of the ecosystem, we can better differentiate how each portion plays an important role in the overall function of the system. We also conclude that more research is needed to understand aquatic productivity of the CRE and how hydrologic dynamics such as freshwater releases to the CRE, local tidal basin run-off and groundwater seepage influence aquatic productivity and ecosystem function. Further, more specific investigation is needed to address the source and role of nutrients to the CRE and contrast the function of riverine-estuarine ecosystems in response to upstream pressures.

## Acknowledgements

We would like to thank Eric Millbrandt, South Florida Water Management District and the Sanibel-Captiva Conservation Foundation staff for field and analytical support. The River, Estuary and Coastal Observing Network is partially supported by NOAA award to E.C.M. and B. Kirkpatrick for the Gulf of Mexico Coastal Ocean Observing System (GCOOS) (#NA16NOS0120018) and the LAT Foundation. The authors would like to thank Peter Doering, John Kominoski, the anonymous reviewers and editor(s) for their efforts and constructive review of this manuscript.

